# Convolutional neural networks trained with a developmental sequence of blurry to clear images reveal core differences between face and object processing

**DOI:** 10.1101/2021.05.25.444835

**Authors:** Hojin Jang, Frank Tong

## Abstract

Although convolutional neural networks (CNNs) provide a promising model for understanding human vision, most CNNs lack robustness to challenging viewing conditions such as image blur, whereas human vision is much more reliable. Might robustness to blur be attributable to vision during infancy, given that acuity is initially poor but improves considerably over the first several months of life? Here, we evaluated the potential consequences of such early experiences by training CNN models on face and object recognition tasks while gradually reducing the amount of blur applied to the training images. For CNNs trained on blurry to clear faces, we observed sustained robustness to blur, consistent with a recent report by Vogelsang and colleagues (2018). By contrast, CNNs trained with blurry to clear objects failed to retain robustness to blur. Further analyses revealed that the spatial frequency tuning of the two CNNs was profoundly different. The blurry to clear face-trained network successfully retained a preference for low spatial frequencies, whereas the blurry to clear object-trained CNN exhibited a progressive shift toward higher spatial frequencies. Our findings provide novel computational evidence showing how face recognition, unlike object recognition, allows for more holistic processing. Moreover, our results suggest that blurry vision during infancy is insufficient to account for the robustness of adult vision to blurry objects.

## Introduction

Humans are remarkably good at recognizing objects across a wide range of viewing conditions, even those that lead to degraded image quality. In particular, blur is a problem that people encounter when viewing most any real-world scene. Any object that lies much closer or further in depth, relative to the point at which the eyes are fixated and accommodated, will appear blurred on the back of the retina (Sprague, Cooper, Reissier, Yellapragada, & Banks, 2016). Blur leads to a loss of high spatial frequency information, such that only the more coarse, lower spatial frequency aspects of the object are registered. A similar loss of high resolution information occurs for objects that appear in peripheral vision, due to retinal and cortical scaling factors that vary as a function of eccentricity (Duncan & Boynton, 2003; Strasburger, Rentschler, & Jüttner, 2011; Virsu & Rovamo, 1979).

Although blur leads to loss of fine detail, behavioral studies have shown that people can still recognize faces and objects after they have been blurred to a considerable extent (Kwon & Legge, 2011). Research has likewise shown that human observers can detect and identify objects at far eccentricities, even beyond 50° (Boucart et al., 2016; Thorpe, Gegenfurtner, Fabre‐Thorpe, & Bülthoff, 2001), although admittedly, foveal recognition is far more accurate. Our ability to recognize blurry or low-resolution objects is crucial for successfully navigating the environment. For example, while driving, it is critical that we can successfully detect unexpected obstacles, traffic signs or pedestrians crossing the street, especially on a rainy day.

To understand how people recognize objects degraded by image blur, it may be useful to consider the performance of deep neural networks, which are believed to provide the most promising current models of the human visual system (Kriegeskorte, 2015; Yamins & DiCarlo, 2016). In particular, convolutional neural network (CNN) architectures (Fukushima, 1980), which apply a series of filtering and pooling operations to mimic the computations of visual cortex (Hubel & Wiesel, 1962), have proven very successful in tasks of object classification (He, Zhang, Ren, & Sun, 2015; Krizhevsky, Sutskever, & Hinton, 2012; Simonyan & Zisserman, 2014) and face identification (Phillips et al., 2018; Taigman, Yang, Ranzato, & Wolf, 2014). Some researchers have even claimed that CNNs have achieved or surpassed human-level performance at tasks of object classification (He et al., 2015; LeCun, Bengio, & Hinton, 2015). It is particularly striking that CNNs, trained on object classification tasks, ultimately learn object representations that are quite similar to those found in the visual cortex of human and non-human primates (Güçlü & van Gerven, 2015; Horikawa & Kamitani, 2017; Khaligh-Razavi & Kriegeskorte, 2014; Yamins et al., 2014). Such neuroscientific findings provide support for the notion that CNNs may perform visual computations that quite closely resemble the feedforward computations of the human visual system.

However, other lines of evidence suggest that CNNs are unusually brittle and lack the robustness of human vision, as modest changes in image quality can sometimes lead to catastrophic failure. For example, CNNs can be severely disrupted if a small amount of adversarial noise (Goodfellow, Shlens, & Szegedy, 2014) or a moderate amount of random Gaussian noise (Dodge & Karam, 2017; Geirhos et al., 2018; H Jang, McCormack, & Tong, 2017) is added to the test image. Image blur has likewise been found to impair CNN performance to a degree that would not be expected for human performance (Dodge & Karam, 2017; Geirhos et al., 2018). In general, it appears that CNNs excel on visual tasks in which test images are drawn from a common statistical distribution as the training images. However, when the test images fall outside of those encountered during training, performance can become severely impaired.

It remains an open question as to whether human vision is more robust than CNN performance because we have acquired a greater range of visual experiences or because the human brain processes visual information in a qualitatively different manner. In this study, we focused on the role of experience, which can be readily modified for CNNs, and asked what types of training experiences might lead to robustness to blur.

We were particularly motivated by the fact that early human visual experience is defined by poor acuity at birth (approximately 20/400) but gradually improves to near-adult levels within the first year of life (Dobson & Teller, 1978; Norcia & Tyler, 1985). The blurry vision of young infants is due to multiple contributing factors, including hypermetropic eyes due to short axial length (Wood, Hodi, & Morgan, 1995), poor accommodation, undeveloped fovea formation, as well as unmyelinated and immature subcortical/cortical circuits (Braddick & Atkinson, 2011; Kiorpes & Movshon, 2004). Although blurry vision at birth may slow the rate of visual learning, it has been posited that this gradual shift from blurry to clear vision might be necessary for the development of more integrative visual processing. This view has received support from studies of patients who had dense cataracts at birth and were subsequently treated at an age of 2-12 months. Since visual acuity was fully corrected by artificial lenses after the treatment, these patients lacked early experience with blurry vision. Despite having many years of compensatory visual experience thereafter, these patients commonly report difficulties with recognizing faces and are severely impaired at detecting changes to the configuration of a face, even though they can still discriminate changes to the local facial features (Geldart, Mondloch, Maurer, De Schonen, & Brent, 2002; Le Grand, Mondloch, Maurer, & Brent, 2001, 2004). Other studies have shown that patients with congenital cataracts are impaired at perceiving the global form of glass patterns (Lewis et al., 2002), suggesting that spatially integrative processing is generally impaired in these patients as well.

A recent study investigated the potential impact of early experience with blurry faces by training a CNN initially with blurry face images, followed by progressively clearer images, and found that the CNN retained considerable robustness to blur after this sequence of training (Vogelsang et al., 2018). In comparison, a CNN trained on faces that progressed from clear to blurry showed good performance for blurry faces but poor performance for clear faces. These findings provide computational evidence in favor of the notion that training with a progression from blurry to clear images may be beneficial for more integrative spatial processing of faces.

The goal of our study was to assess the generality of these findings by assessing whether the processing of blurry objects would necessarily benefit from a sequence of blurry to clear image training. On one hand, it seems natural to expect that a common training regime should lead to the same benefits for the processing of non-face objects. On the other hand, the tasks of face recognition and object recognition are quite distinct and present different computational challenges to the visual system. Face recognition is a challenging task that relies on dedicated neural circuitry to distinguish subtle differences between stimuli that share a common higher order structure (Kanwisher, McDermott, & Chun, 1997; Tong, Nakayama, Moscovitch, Weinrib, & Kanwisher, 2000) by relying on norm-based coding (Chang & Tsao, 2017; Freiwald, Tsao, & Livingstone, 2009; Loffler, Yourganov, Wilkinson, & Wilson, 2005). By contrast, object recognition requires distinguishing among a much more heterogeneous set of stimuli that can vary along a wide array of features and object dimensions (Hebart, Zheng, Pereira, & Baker, 2020). As a consequence, a much more diverse set of diagnostic features must be learned in tasks of object recognition. Such differences in the visual stimuli and associated task demands may account for why face recognition relies more heavily on configural and holistic processing (Le Grand et al., 2004; Tanaka & Simonyi, 2016).

There is also research to suggest that face recognition relies on lower spatial frequency information when compared to the subordinate-level discrimination of certain non-face object categories, such as airplanes (Harel & Bentin, 2009). Although face recognition tends to be most accurate for stimuli presented in a mid-range of spatial frequencies with peak performance centered around 8 cycles per face stimulus (Gaspar, Sekuler, & Bennett, 2008; Näsänen, 1999; Peli, Lee, Trempe, & Buzney, 1994), recognition remains possible for faces that have been low-pass filtered to a cut-off frequency of only 2-3 cycles per stimulus (Kwon & Legge, 2011). Moreover, low-pass filtered faces have been found to engage holistic processing, whereas high-pass filtered faces do not (Goffaux & Rossion, 2006). Taken together, these findings raise the possibility that blur applied to faces and non-face objects may lead to differential effects on visual processing.

In this study, we trained CNNs on a progressive sequence of blurry to clear images, and evaluated whether robustness was retained at the end of training for face- and object-trained networks. We found that the face-trained CNN successfully maintained robustness to blur at end of training; moreover, during intermediate training stages it could successfully generalize across both blurry and clear faces. By contrast, object-trained CNNs showed elevated performance for the most recently trained blur level but poor generalization to other blur levels. By the end of training, the CNN was no longer robust to blurry objects.

We devised an analysis to characterize the spatial frequency tuning of the face- and object-trained networks, not only for the first convolutional layer but also for higher layers. This analysis revealed dramatic differences between the two networks. The face-trained network successfully maintained a preference for lower spatial frequencies, whereas the object-trained network showed a progressive shift towards higher spatial frequencies after each stage of training. As a control, we trained a CNN on object images that were either low-pass filtered using a Gaussian kernel or matched to the Fourier power spectrum of the face images. Even after these image manipulations, followed by our procedure for blurry to clear training, the CNNs were unable to retain robustness to blur. We performed additional controls with CNNs trained on subordinate-level categorization of dogs or birds as well as CNNs trained on superordinate-level categorization of animate versus inanimate objects. In all cases, blurry to clear training with these object stimuli proved unsuccessful at conferring robustness to blur.

Our results suggest that more holistic, lower spatial frequency representations can be used to support successful recognition of faces across a range of blur levels. By contrast, object recognition requires higher spatial frequency information to achieve optimal performance. With a progression of blurry to clear training images, the object-trained CNN learns to leverage this new, more discriminating information, modifying its weights to such a degree that it no longer successfully processes object images presented at an earlier trained blur level. Although early infant experiences with blur may help encourage the development of more holistic face processing, our findings with object-trained CNNs suggest that these early experiences are insufficient to account for people’s ability to recognize blurry objects. More generally, this study provides novel computational evidence that demonstrates how the spatial properties of faces and objects are profoundly different, and can lead to different visual learning strategies by recognition systems.

## Methods

### Visual stimuli

Face images were collected from the FaceScrub database (Ng & Winkler, 2014), which consisted of 100,000 face images sampled from 530 celebrities. The dataset only provided the URLs to the images, and if image URLs were invalid (as of October 13, 2019), those images were excluded. We also excluded any face identities with fewer than 100 examples. This resulted in a final face image dataset with 395 face identities to train CNNs on face recognition. Object images were obtained from the ILSVRC-2012 or ImageNet database (Russakovsky et al., 2015), which has 1,000 object categories with roughly 1.25 million training and validation images. All 1,000 object categories were used to train the object-trained CNNs. All stimuli were converted to grayscale and resized to 224 × 224 pixels to meet the image processing requirements for CNNs.

For our behavioral face recognition task, we chose 10 celebrities (5 females and 5 males) who we considered likely to be well known to the general public: Jennifer Aniston, Mila Kunis, Ellen Degeneres, Selena Gomez, Anne Hathaway, Jim Carrey, Matt Damon, Robert Downey Jr., Ryan Gosling, and Samuel L. Jackson. One of the authors reviewed and sorted out mislabeled or idiosyncratic photographs of faces. Furthermore, we excluded any images that shared a pixel-wise correlation exceeding 0.9 with any other image. The final face image set consisted of 80 images per celebrity or 800 images in total. Regarding image variability, the face images of a given celebrity could vary to a considerable degree due to variations in lighting, viewpoint (ranging from front to three-quarter view), facial expression, hairstyle, make-up, facial hair, age and/or accessories worn (e.g., glasses, hat). We applied a Gabor wavelet pyramid model with 5 spatial scales and 8 orientations to calculate the Pearson correlational similarity of simulated complex cell responses to the images. Normalization was first applied to the all responses at a given spatial scale to control for greater power at lower spatial frequencies. The pairwise correlational similarity of face images was somewhat greater for within-celebrity comparisons (mean r = 0.464, sd = 0.141) than between celebrities (mean r = 0.405, sd = 0.122).

For the behavioral object recognition task, 16 object categories were selected to compare human and CNN performance: bear, bison, elephant, hamster, hare, lion, owl, tabby cat, airliner, couch, jeep, schooner, speedboat, sports car, table lamp, and teapot. Half of the object stimuli were animate and the other half were inanimate. Fifty images per category from the ImageNet validation dataset were used, and thus we had 800 images in total. We performed the same Gabor wavelet pyramid model analysis to the object images. The correlational similarity of the object images was somewhat greater for within-category comparisons (mean r = 0.292, sd = 0.159) than between category (mean r = 0.255, sd = 0.148). As expected, the object images were more heterogeneous than the face images, and within-category (or within-identity) images shared somewhat greater low-level similarity than between-category images.

To generate the blurred images, we applied a Gaussian kernel to each image, adjusting the standard deviation (σ) of the Gaussian function to attain different levels of blur. All image processing was performed using MATLAB. For both behavioral experiments, all images were upsampled by a factor of 2 for presentation on a CRT monitor at a size of 19 × 19 degrees of visual angle.

### Participants

We recruited 20 participants to take part in the behavioral object recognition study. A separate group of 20 participants were recruited to take part in the face recognition study. Each of the two studies required approximately 1 hour to complete. All participants reported having normal or corrected-to-normal visual acuity and provided informed written consent. The study was approved by the Institutional Review Board of Vanderbilt University. Participants were compensated monetarily or through course credit.

### Behavioral experiments

We measured the abilities of human observers at recognizing faces and objects presented with varying degrees of blur (σ = 0, 1, 2, 4, 8, 12, 16, 20, 24, and 32). Here, σ = 0 indicated clear images without any blurring. Eight face images per celebrity were assigned to each blur level for the face recognition task, while five images per object category were assigned to each blur level for the object recognition task. Both experiments consisted of a total of 800 images, with 80 images presented at each blur level. Image assignment across blur levels was counterbalanced across participants and the order of image presentation was randomized.

Each visual stimulus was briefly presented for 200 ms on a gray background, subtending a visual angle of 19°. After stimulus presentation, participants were asked to report the face identity or object category by entering in a corresponding number code on a numerical pad. The correspondence between the number code and the stimulus identity remained on the screen throughout the study. The mapping between number codes and stimulus identity was counterbalanced across participants. The experiment required approximately 1 hour to complete, including informed consent, instructions, and debriefing. The experiment was implemented using MATLAB and the Psychophysics Toolbox (http://psychtoolbox.org/).

### Training of convolutional neural networks

The majority of all CNN experiments and analyses were performed using AlexNet, which can achieve a high level of classification performance while still being quite fast to train from scratch (Krizhevsky et al., 2012). We performed supplementary analyses using VGG-19, which is a deeper CNN with greater learning capacity (Simonyan & Zisserman, 2014).

With the face dataset of 395 celebrities, we divided the images into separate training and validation sets using an approximately 90/10 split. On average, this led to 117 examples per identity for training and 13 examples per identity for validation. For object images obtained from ImageNet, we used their training images (~1.2 million) for training and their validation dataset (50k images) for testing the CNNs. For data augmentation, the training images were randomly rotated from −10° to +10° and about half were flipped about the vertical axis. Across all images within a training set, we calculated the mean and standard deviation of the pixel intensity values and used these values to normalize the pixel intensities of the images.

The models were trained using stochastic gradient descent with a fixed learning rate of 0.01, momentum of 0.9, and weight decay of 0.0001. To train the network initially with blurred images, we applied a Gaussian kernel with the standard deviation of σ = 8, and subsequently reduced the blur level to 4, 2, 1, and 0. The blur level was changed every 100 training epochs for the face recognition task and every 10 training epochs for the object recognition task. Given that the number of training examples per category of ILSVRC-2012 was approximately 10 times larger than that of FaceScrub, the networks in both tasks processed similar numbers of training images per category for each blur level. For comparison, a control CNN was trained with only clear images using the same number of training epochs. All training procedures were implemented in PyTorch on a workstation equipped with multiple GPUs.

### Receptive field analysis

We fitted a 2D elliptical Gabor model to the first-layer receptive fields (11 × 11 pixels) of the trained CNNs. The function we used obtained the best fitting model after sampling from 100 different starting points using a gradient descent method. The filters whose R-squared value was less than 0.4 were excluded from analysis. After fitting, the average of standard deviation values of the 2D Gaussian envelope was determined as the size of the receptive field.

### Peak spatial frequency in tuning curves

To estimate the peak spatial frequencies of the network across layers, we devised a method in which we presented grating patterns to CNNs and examined the responses of feature maps across layers. Specifically, the gratings were created by sinusoidal patterns using 15 orientations (0, 12, ..., 168 in degree), 25 spatial frequencies (4.48, 8.96, ..., 112 cycles/stimulus), and 4 phases (0, 45, 90, 135 in degree). We measured the average responses to the gratings from individual feature maps across convolutional layers and plotted the tuning curves for spatial frequency by averaging across orientations and phases. Each tuning curve was normalized to a range from 0 to 1. The peak spatial frequency was determined from each tuning curve as it yielded the maximum. This peak spatial frequency indicated which spatial frequency was mostly preferred by each feature map of the network.

### Spatial frequency control images

As a supplementary analysis, we manipulated the spatial frequency content of the training object images in two ways. First, we calculated the average amplitude spectrum of all training face images and replaced the amplitude spectrum of individual training object images with the average amplitude spectrum from the faces. This was done by performing the fast Fourier transform on each object image in MATLAB, adjusting the amplitude spectrum accordingly, and then performing the inverse fast Fourier transform to reconstruct the amplitude-matched object image. Our second approach relied on low-pass filtering applied to training object images using a cutoff frequency of either 32 or 16 cycles per image. This was done by zeroing out all amplitude values below the cutoff frequency in the Fourier domain, and then performing the inverse fast Fourier transform to reconstruct the image.

### Code and data availability

The experimental code, PyTorch code, and human behavioral data will be made available on open science framework upon publication of this work. Visual stimuli can be obtained via the FaceScrub and ImageNet databases. https://osf.io/95pek/?view_only=7a634add763c4e2a81c2fe1830df23d6

## Results

We first evaluated the performance of human observers and CNNs on their ability to recognize faces or objects presented with varying levels of blur. We selected 10 celebrities from the FaceScrub dataset (Ng & Winkler, 2014) and 16 object categories (8 animate, 8 inanimate) from the ImageNet dataset (Russakovsky et al., 2015) for these forced-choice recognition tasks. To quantify CNN performance, we trained a separate model of AlexNet on either faces or objects, using clear grayscale versions of all of the training images from the corresponding data set, and then evaluated test performance using independent test images (see **Methods**).

**Figure 1** shows performance accuracy plotted as a function of blur level, as indicated by the standard deviation (σ) of the Gaussian kernel used for filtering. Human and CNN performance was about equally accurate for clear images (σ = 0) but quickly diverged at modest levels of blur. Although CNN performance never quite declined to chance level (1/10 or 10%) for faces (**Figure 1A**), performance became very poor at a blur level of 8 and approached near floor performance by a blur level of 12. In comparison, human performance remained above 50% accuracy at a blur level of 12. CNNs were particularly vulnerable to blurry objects with performance falling to near-chance levels at a blur level of 4, whereas human performance remained significantly above chance at a blur level of 16 (**Figure 1B**). It can also be seen that blurry objects were more challenging to recognize than blurry faces for both CNNs and human observers.

**Figure 1.**
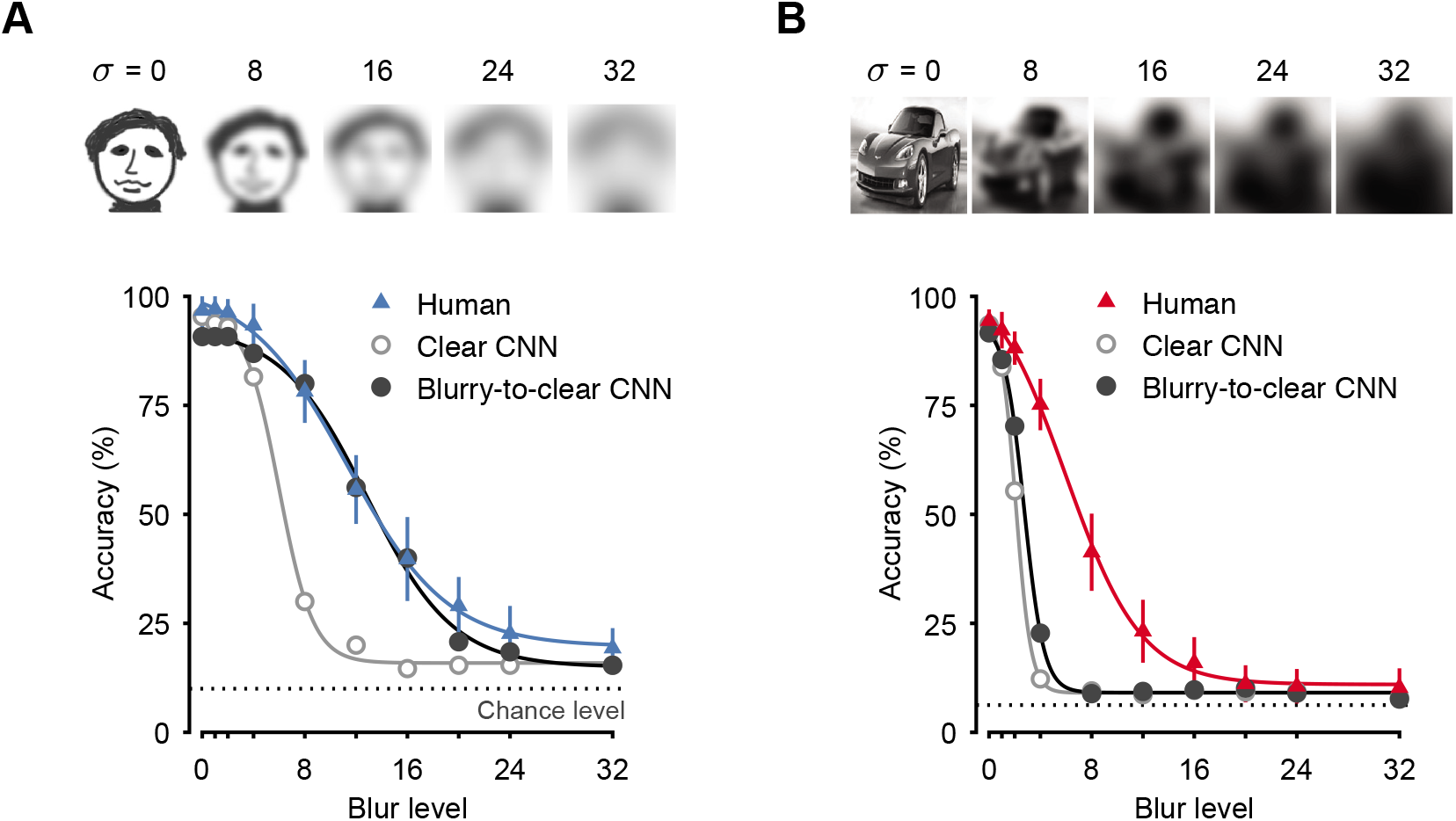
**A** Face recognition accuracy of human observers (blue triangles), AlexNet trained on clear face images (gray open circles), and AlexNet trained on a sequence of blurry to clear faces (black filled circles) when tested with a wide range of blur levels (chance level performance with a dashed line, 1/10 or 10%). The solid lines represent a logistic function fitted to the data. Note that cartoon face images are shown to illustrate the amount of blur instead of the examples of celebrities due to potential copyright issues. **B** Object recognition accuracy of human observers (red triangles), AlexNet trained on clear objects (gray open circles), and AlexNet trained on a sequence of blurry to clear objects (black filled circles) when tested with a wide range of blur levels (chance level performance, 1/16 or 6.25%). Error bars indicate ±1 standard error of the mean.

We sought to determine whether the training of CNNs on a sequence of blurry to clear images would lead to enhanced robustness to blur. CNNs with initially randomized weights were trained on either faces or objects with decreasing levels of blur applied to the images across successive stages of training (σ = 8, 4, 2, 1 or 0). After the final stage of training, we evaluated the performance of these CNNs with the same subset of blurry faces and objects that were used to assess human performance.

**Figure 1** reveals a striking divergence in performance between the face-trained and object-trained CNNs. The blurry to clear face-trained CNN performed much better at recognizing blurry faces than the clear-trained CNN; moreover, it provided a good approximation of human performance, which is quite robust to blur. Performance was even improved at blur levels that extended beyond those that were encountered during blurry to clear face training (σ = 12 and 16). By contrast, the blurry to clear object-trained CNN performed hardly better than its clear-trained counterpart, and both were far less robust to object blur than human observers. These findings suggest that blurry to clear trained CNNs may provide a suitable model to account for the face recognition performance of human observers but not their object recognition performance.

How does the performance of these CNNs change across successive stages of training? **Figure 2A** shows the performance accuracy of the face-trained CNN, evaluated after each stage of training (blue curve) on test images from 395 celebrities presented with varying levels of blur. After the first stage of training (σ = 8, leftmost plot), the CNN performed well at the highest blur level but was unable to generalize to images with little to no blur (σ ranging from 0-2) that had yet to be trained. However, after the second stage of training (σ = 4), the face-trained CNN performed well across the full range of blur levels, generalizing well even to clear images that had yet to be encountered in training. Subsequent stages of training led to modest improvement for clear faces and faces with minimal blur, accompanied by a modest reduction in test performance at the highest blur level (σ = 8). By the end of training (rightmost plot), the CNN exhibited a modest advantage for clear images yet remained robust across the full range of previously trained blur levels. These findings are largely consistent with those reported by Vogelsang et al. (2018).

**Figure 2.**
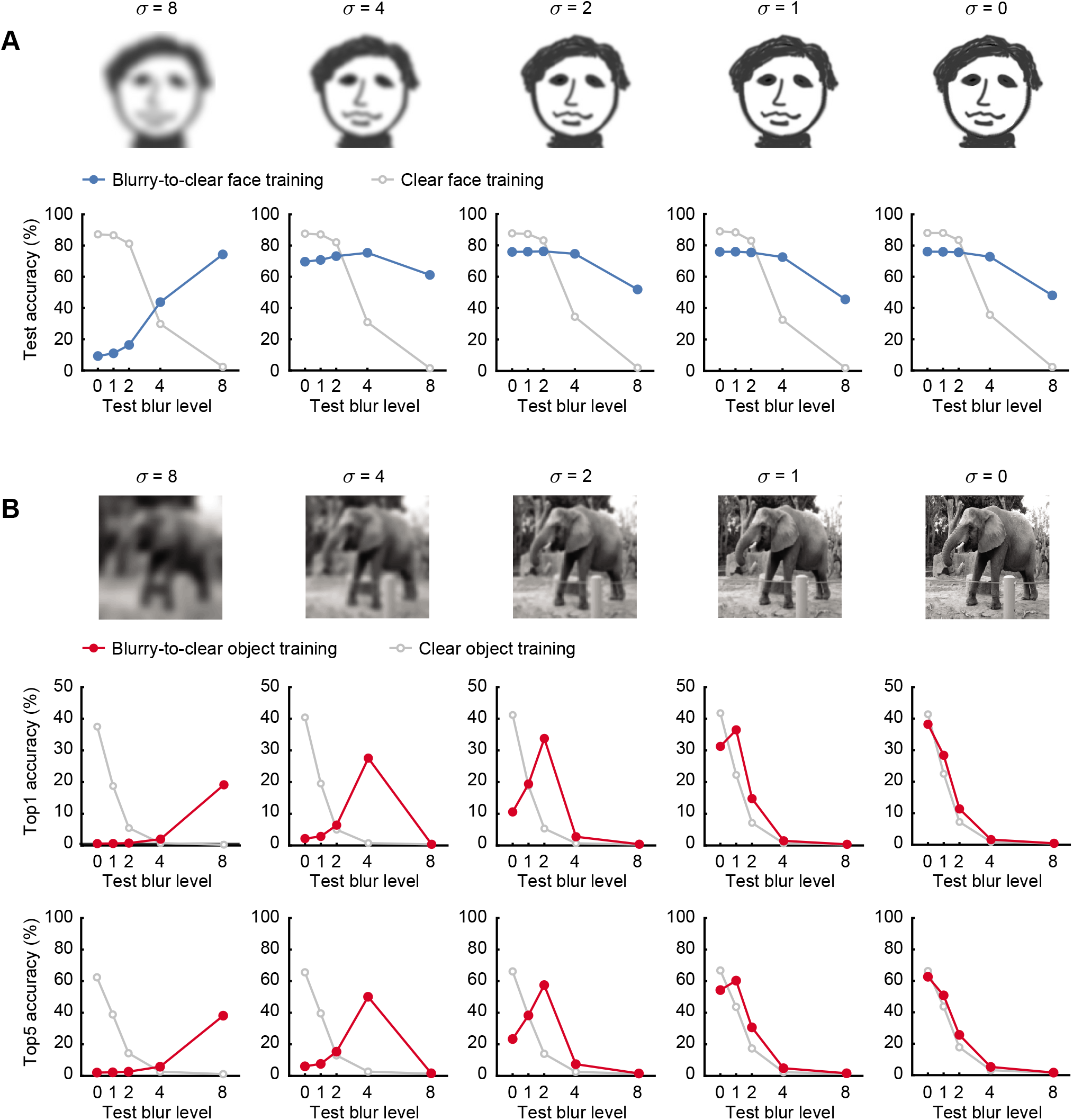
Performance accuracy of AlexNet after training with faces (**A**) or objects (**B**) presented at different blur levels (σ = 8, 4, 2, 1 or 0) over a series of training stages. Gray curves indicate the performance of control CNNs trained on clear images only. To illustrate the amount of blur applied at each training stage, a cartoon face at each blur level is shown instead of the examples of celebrities to avoid copyright issues.

For comparison, we trained a control CNN with the same number of training images but used clear faces only. The control CNN (gray curve) exhibited excellent performance with clear and minimally blurred images (σ ranging from 0-2) after only a single stage of training, outperforming the blurry to clear face-trained in this range. However, the control CNN performed poorly with faces presented at high blur levels of 4 or 8, indicating poor robustness to blur. Subsequent stages of training led to negligible changes in performance.

We observed a very different pattern of results for the CNN trained on a sequence of blurry to clear object images, which was trained using 1000 categories of objects obtained from ImageNet. Although the first stage of training with blurred objects led to improved performance at that blur level with top1 accuracy of 19.2% (**Figure 2B**, σ = 8, top row, leftmost plot), training at the next blur level (σ = 4) led to a loss of robustness to the originally trained level of blur, with performance plummeting to an accuracy of 0.37% (chance level, 0.1%). Subsequent stages of training led to similar declines in robustness to previously trained levels of blur. After the final stage of training with clear images (σ = 0), performance across blur levels was very similar to that of the control CNN trained exclusively on clear objects (gray curve, **Figure 2B**, rightmost plot). We also calculated the top5 accuracy of the object-trained CNNs to ensure that their lower performance accuracy would not obscure our ability to find evidence of successful generalization across changes in blur level. Here, responses were scored as correct if the correct category was among the 5 highest softmax responses of the CNN. We found that top5 accuracy (**Figure 2B**, bottom row) revealed the same pattern of results as top1 accuracy.

One potential concern with respect to the blurry to clear object training results might be that the learning capacity of AlexNet was insufficient to maintain the knowledge acquired from earlier stages of training. We conducted the same experiment on a higher capacity CNN, VGG-19 (Simonyan & Zisserman, 2014), to address this issue and observed essentially the same pattern of results for both faces and objects (**Figure 3**). Our findings suggest that training with a sequence of blurry to clear objects is insufficient for CNNs to learn stable object representations that remain robust to blur.

**Figure 3.**
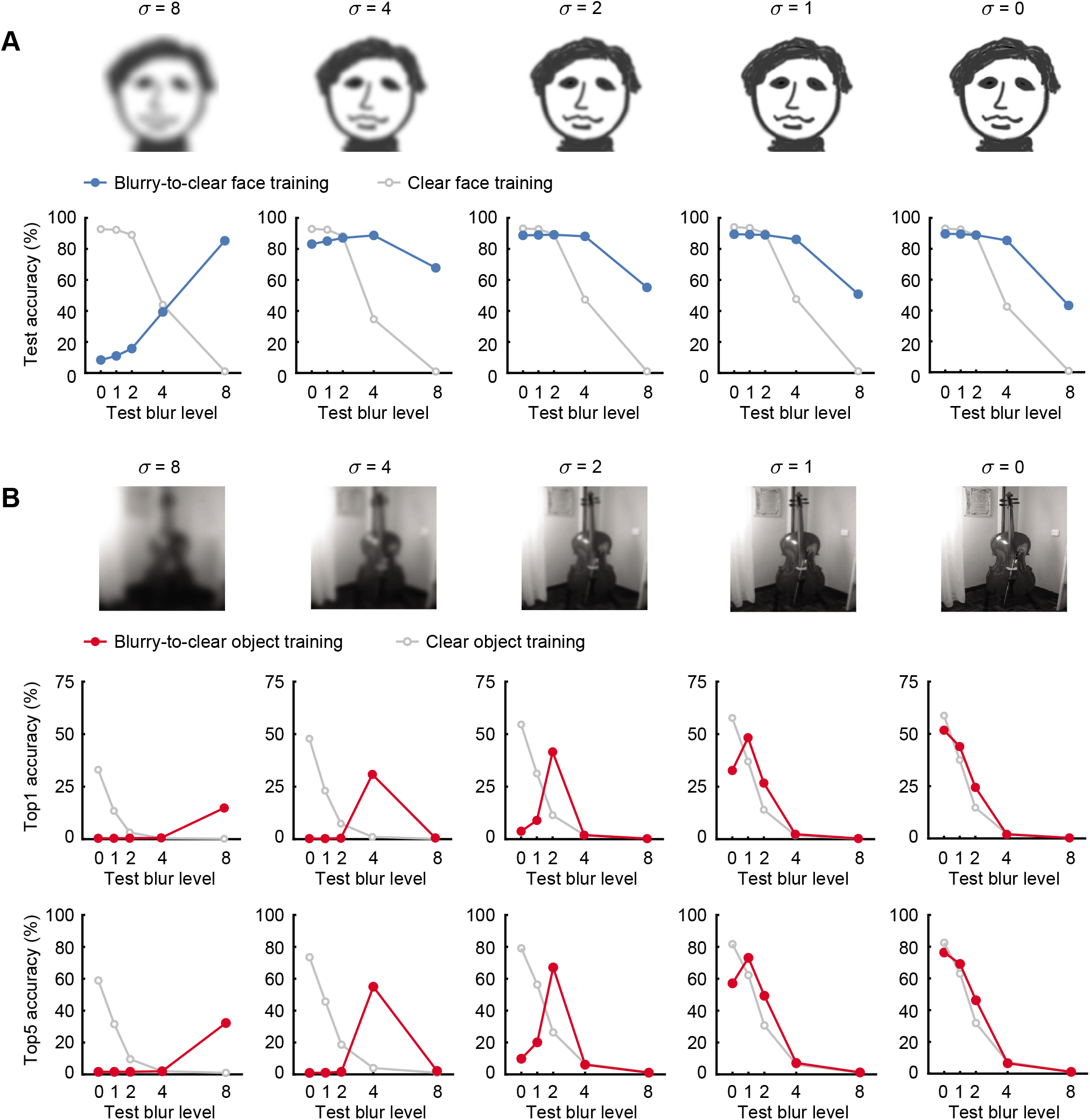
Performance accuracy of VGG-19 after training with faces (**A**) or objects (**B**), following the conventions described in **Figure 2**. Cartoon face images are shown at each blur level due to potential copyright issues.

To understand how blur training with faces or objects modified the representations learned by the CNNs, we visualized the receptive fields learned by the first convolutional layer. Each row in **Figure 4A** depicts how the receptive field of an example unit changes across successive stages of training. For the face-trained network, the receptive fields remained quite stable across training stages, with a preference for large, coarse features. By contrast, the CNN trained on blurry to clear objects exhibited marked changes across training periods — the receptive fields appeared to shrink in size and shifted towards preferring higher spatial frequencies. These findings indicate that training with progressively clearer objects induced the CNN to learn finer spatial representations.

**Figure 4.**
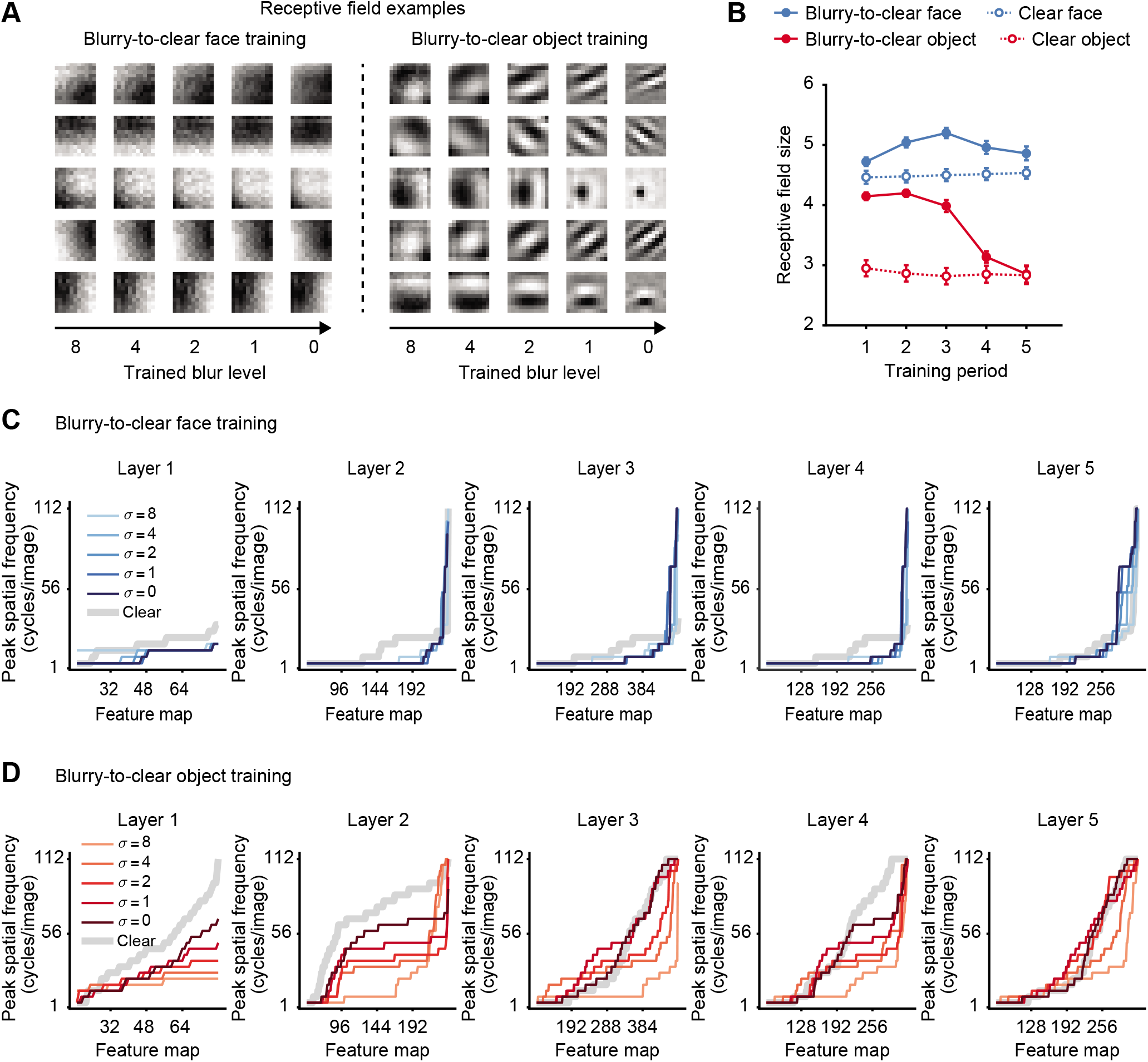
**A** Examples of the learned receptive fields obtained from a CNN trained on a sequence of blurry to clear faces (left) or blurry to clear objects (right). **B** Receptive field sizes were measured after each training period for both blurry to clear trained CNNs and CNNs trained on clear images only. **C** Peak spatial frequency preferences of face-trained CNNs across successive stages of training, with separate plots shown for each convolutional layer. For visualization purposes, the feature maps are sorted by their peak spatial frequency preference. The gray lines indicate the peak spatial frequency preferences of the clear face-trained network to serve as a reference. **D** Peak spatial frequency preferences of CNNs trained on object recognition, following the conventions of **C**.

We quantified the receptive field size of these first layer representations by fitting a 2D Gabor model and calculating the standard deviation values corresponding to the Gaussian profile of the fitted model. This analysis revealed that blurry to clear face training allowed the CNN to learn and maintain larger receptive fields than the CNN trained on clear faces only (**Figure 4B**). In comparison, the blurry to clear object-trained CNN had large receptive fields initially but these shrunk in size with training such that by the final stage, they were no larger than those of the clear object-trained CNN. These results are concordant with the pattern of performance accuracy that we observed as a function of blur level after the final stages of training (**Figure 2**).

To understand how the spatial frequency tuning of the CNNs was affected by the training, we calculated how strongly units in each layer responded to sinewave grating patterns presented across a range of spatial frequencies, orientations, and spatial phases (see **Methods**). This analysis allowed us to characterize the spatial frequency preferences not only for layer 1 but for higher layers as well. We calculated the preferred spatial frequency of all feature-tuned units in convolutional layers 1 through 5 of AlexNet.

The plots in **Figure 4** depict the preferred spatial frequency of all feature-tuned units in a given layer, sorted by preference from low to high. Focusing on layer 1 of the blurry to clear face-trained CNN (**Figure 4C**, left), it can be seen that spatial frequency preferences remained stable across training stages (σ from 8 to 0) and were considerably lower than those of the clear face-trained CNN (gray curve). A very similar pattern of results could be seen for the blurry to clear face-trained CNN in layers 2-5. We performed a Mann-Whitney U test to determine whether the population of feature-tuned units in a given layer exhibited significant changes in spatial frequency preference between successive stages of training. The units in layers 1 through 5 all showed a significant change in spatial frequency preference (p < 0.005 in all cases) between the first and second stages of blurry to clear training (σ = 8 vs. σ = 4) but showed no significant changes thereafter. After training was complete, the blurry to clear face-trained CNN preferred lower spatial frequencies than the clear face-trained CNN (p < 0.001 for layers 1-4, p < 0.05 for layer 5). We conclude that the blurry to clear face-trained CNN exhibited strong and stable preferences for lower spatial frequencies in most units throughout the network.

These findings can be contrasted with the blurry to clear object-trained CNN (**Figure 4D**), which exhibited a marked shift in preference toward higher spatial frequencies over the course of training. This trend could be observed in all 5 convolutional layers, and was particularly prominent in layers 2 and 3, which showed a significant shift in preference toward higher spatial frequencies after every stage of training (p < 0.001 in all cases). After the final stage of training, the blurry to clear object-trained CNN did exhibit a preference for somewhat lower spatial frequencies in layers 1 and 2 than did the control CNN (p < 0.001) but such differences were much weaker or negligible in the higher layers (layer 3, p = 0.36; layer 4, p = 0.0054; layer 5, p = 0.95). A direct comparison of the face- and object-trained CNNs (**Figures 4C vs. 4D**) further reveals how the object recognition task led to preferences for a higher as well as much wider distribution of spatial frequencies, when compared to face recognition. A similar analysis of preferred spatial frequency was performed on the VGG-19 networks, which revealed stable preferences for lower spatial frequencies during blurry to clear face training and a shift in preference toward higher spatial frequencies during blurry to clear object training (see **Supplemental Materials**). These findings concur with the notion that faces can be processed in a more holistic manner than objects, though to our knowledge, our computational approach for demonstrating this effect is quite different from previous approaches (Goffaux, Hault, Michel, Vuong, & Rossion, 2005; Le Grand et al., 2004; Richler & Gauthier, 2014; Sinha, 2002; Tanaka & Simonyi, 2016).

Why did the representations for faces change so little as the training images progressed from blurry to clear, while the representations for objects changed so dramatically? **Figure 5** shows a continuous plot of the accuracy of training performance over time for both face and object recognition tasks. The face recognition network was able to achieve excellent performance very quickly at the initial blur level of 8, and by a blur level of 4, performance was near ceiling, indicating that the network had very little left to learn. These findings are consistent with the strong generalization that was observed across all blur levels after the CNN was trained on faces with a blur level of 4 (**Figure 2A**). In comparison, the object recognition network showed slower and more gradual improvement, implying that more remained to be learned in order to adapt to each new blur level, which provided new higher spatial frequency information to leverage for improved performance. This presumably led to overwriting of the representations that supported earlier robustness to blur.

**Figure 5.**
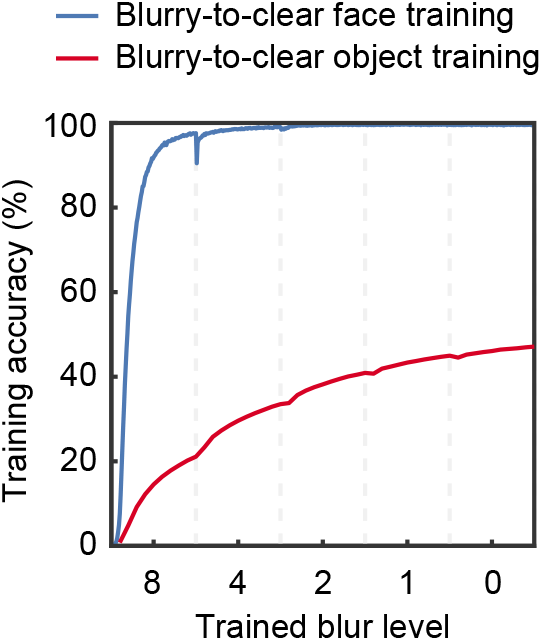
Performance accuracy on training images for CNNs trained on a progression of blurry to clear faces (blue) or objects (red). Vertical dashed lines indicate the transitions between blur levels.

One might ask whether controlling for differences in the overall spatial frequency content of faces and objects might allow CNNs to learn more blur-robust or holistic representations of objects. A Fourier power spectrum analysis indicated that the object images did indeed contain greater power at higher spatial frequencies when compared to the face images (**Figure 6A**). If the object images were matched to the power spectrum of the face images, would blurry to clear object training lead to robustness to blur? We suspected that this would be unlikely, given that the spatial frequency information that is needed for successful classification of face identity or object category might be quite distinct from the range of spatial frequencies that are contained in the face or object images.

**Figure 6.**
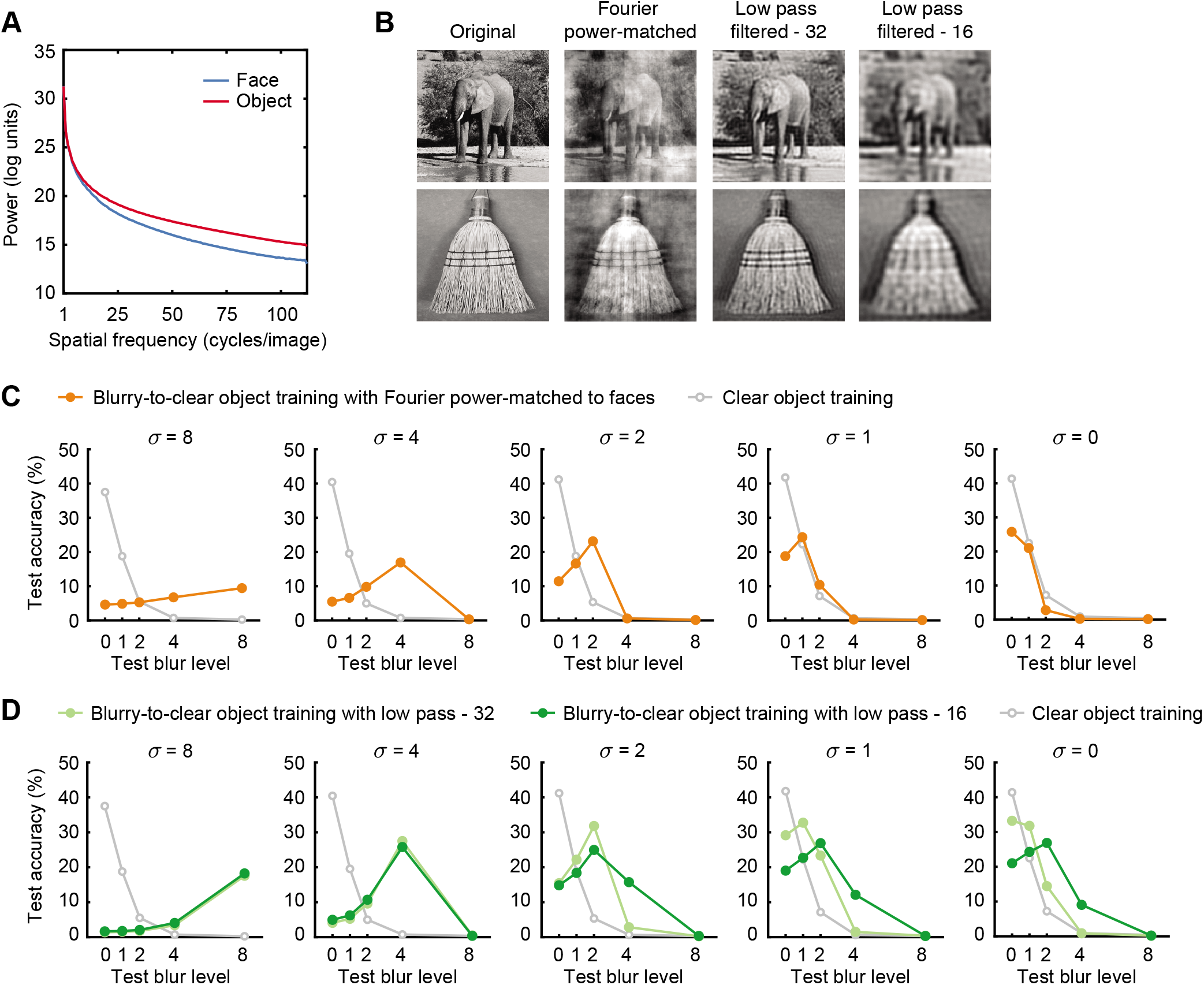
**A** Average power spectrum of face and object training images plotted on a log scale (ordinate) as a linear function of spatial frequency. **B** Examples of original object images (first column), object images with Fourier power spectrum matched to the average power spectrum of the face training dataset (second column), and low-pass filtered images with a cut-off frequency of either 32 cycle/images (third column) or 16 cycle/images (fourth column). **C** Recognition accuracy for a CNN trained on blurry to clear objects, after objects were first matched to the Fourier power spectrum of faces (orange curve). For comparison, performance of the CNN exclusively trained on the original clear objects is also shown (gray). **D** Object recognition accuracy of CNNs trained on blurry to clear objects, after the objects were low-pass filtered with a cut-off frequency of 32 cycles per stimulus (light green) or 16 cycles per stimulus (dark green). Again, the CNN originally trained on clear objects is shown in gray.

Nevertheless, we thought it would be informative to perform this control analysis. We adjusted the set of object images to match the average power spectrum of the training face images (**Figure 6B**), and then trained a CNN with these blurry to clear object images. The trained CNN was then tested with the original set of object test images with varying levels of blur applied. Despite controlling for differences in spectral power during training, blurry to clear object training still failed to lead to robustness to blur (**Figure 6C**), and the pattern of results was almost identical to those observed with the original blurry to clear object-trained CNN (**Figure 2B**). To extend this analysis, we constructed two new sets of object images that were low-pass filtered with a cut-off frequency of either 32 or 16 cycles per stimulus. Again, CNNs trained on these blurry to clear object images failed to retain robustness to blur at the end of training (**Figure 6D**). With more severe low-pass filtering of 16 cycles, the CNN showed some ability to generalize to modest levels of blur but higher levels of blur remained problematic (σ = 4 or 8). Moreover, these modest improvements for blurry objects came with the cost of poorer performance with clear objects. Our findings indicate that the inadequacy of blurry to clear object training for inducing robustness to blur cannot be attributed simply to differences in spatial frequency content between objects and faces or to the presence of high spatial frequency information in objects.

We further sought to determine whether different levels of object categorization might have an impact on the efficacy of blurry to clear image training. Face recognition requires performing fine-grained subordinate-level discrimination between visually similar stimuli that share a common configuration across different identities, thereby presenting challenges that differ from basic-level object categorization (Diamond & Carey, 1986; Gauthier & Tarr, 1997). Therefore, it is conceivable that a CNN trained exclusively on subordinate-level object categorization might benefit from blurry to clear image training. On the other hand, developmental research has suggested that infants progress in their visual categorization skills by first learning about superordinate categories (e.g., animate vs. inanimate), followed by basic-level categories, and then subordinate-level categories (Mandler & McDonough, 1993; Quinn, 2004). This led us to also consider whether blurry to clear image training might enhance the robustness of superordinate categorization.

To investigate subordinate-level object categorization, we leveraged the semantic WordNet hierarchy of ImageNet to select 116 different dog breeds and 52 bird species. Separate CNNs were trained to classify either dogs or birds using the same training procedures as before. Our analyses revealed that blurry to clear image training on these subordinate-level categorization tasks failed to induce sustained robustness to blur, as performance was very poor at high levels of blur (**Figure 7a**, rightmost panel). For the dog-trained CNN, we observed a modest improvement after blurry to clear training for images with moderate blur (σ = 2) but this came with the associated cost of poorer performance for clear images (σ = 0) when compared to the clear-trained CNN. The bird-trained CNNs exhibited lower accuracy overall, and blurry to clear image training led to poorer performance for clear images while failing to improve performance for blurry images.

**Figure 7.**
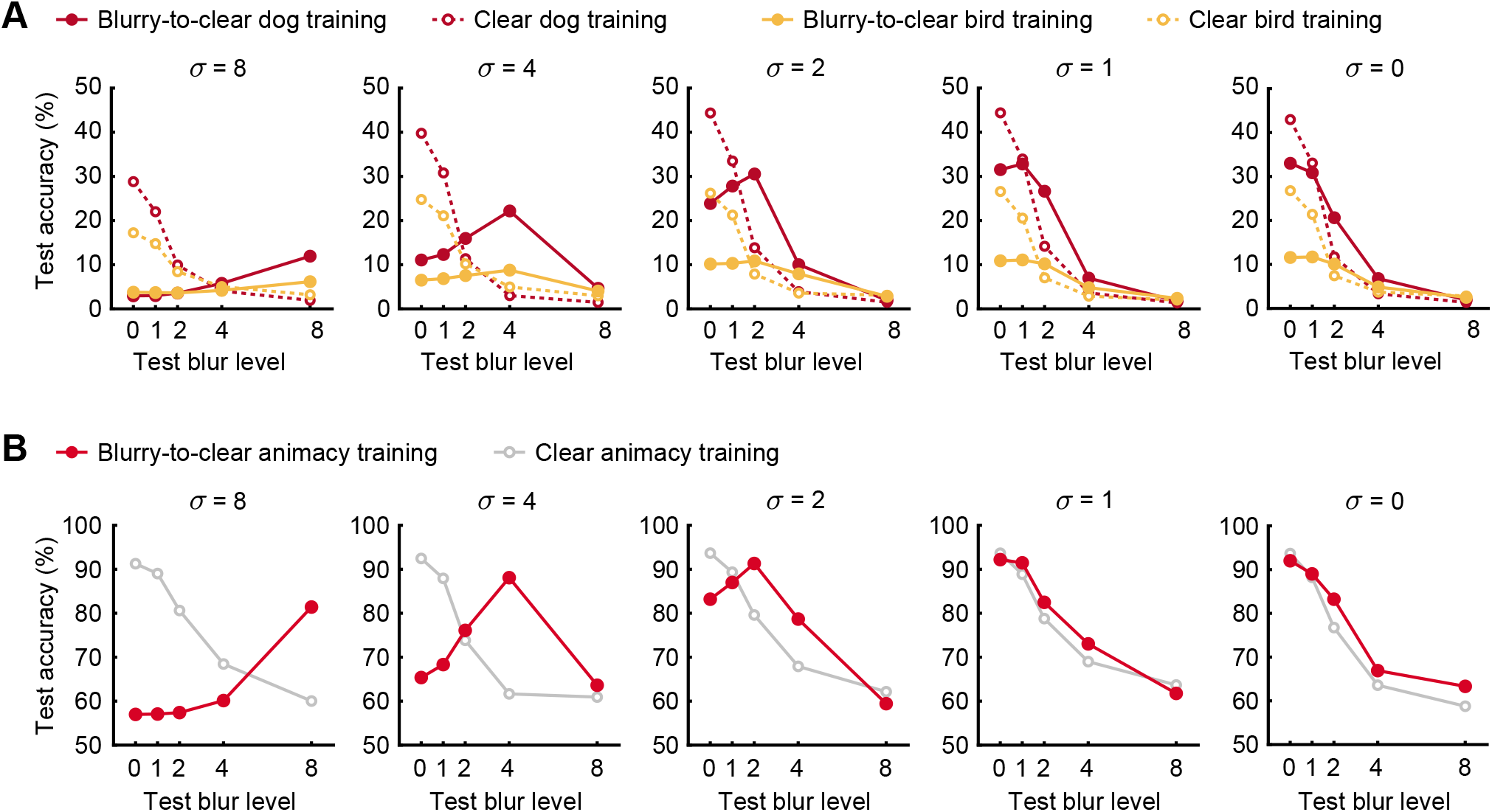
**A** Comparison of CNNs trained on blurry-to-clear (solid) or clear (dashed) images of different dog breeds (red) or bird species (orange). **B** Comparison of CNNs trained on blurry-to-clear (red) or clear (gray) images of ImageNet objects to perform an animate/inanimate discrimination task.

We also investigated superordinate-level classification by sorting the ImageNet categories according to 407 animate categories and 522 inanimate categories. When a CNN was trained to perform animate/inanimate classification using a sequence of blurry to clear images, it improved at the currently trained blur level but lost robustness to the previously trained blur level (**Figure 7b**). After training was completed, the blurry to clear trained CNN exhibited performance that scarcely differed from the clear-trained CNN. Collectively, these results demonstrate that CNNs trained on blurry to clear objects fail to maintain robustness to blur, regardless of whether they are trained on superordinate, basic-level or subordinate-level categorization. By contrast, face-trained CNNs, which must learn to distinguish faces at the subordinate level, can readily acquire robustness to blur from blurry to clear image training.

## Discussion

In this study, we investigated whether CNNs trained with a developmentally motivated sequence of blurry to clear images would be able to attain robustness to blur that could better match human performance. Training with blurry to clear faces allowed CNNs to achieve considerable robustness to blur, consistent with a previous report by Vogelsang et al. (2018). Moreover, our blurry to clear face-trained CNN provided a very good approximation of human recognition accuracy for blurry faces (**Figure 1A**), suggesting that such trained networks may provide a more suitable model of human face processing. It was noteworthy that at an intermediate stage of training, the CNN trained with a modest level of blur (σ = 4) could successfully generalize across both blurry and clear faces. These findings demonstrate that sufficient information was present in the lower spatial frequency range of these face stimuli to allow for successful learning and generalization. We characterized the spatial frequency tuning preferences of this face-trained network across successive stages of training, and found that the CNN was able to learn stable lower spatial frequency representations of the face stimuli and support successful recognition across a range of blur levels.

However, it should be emphasized that the learning of these lower spatial frequency representations of faces was not a foregone conclusion. CNNs that were trained exclusively with clear faces showed poor generalization to blurry face images and exhibited a preference for higher spatial frequencies. These results are consistent with the proposal that visual experience with blurry faces in early infancy may be critical for the development of holistic face processing and that a lack of such early experiences with blurry faces may bias recognition to favor finer spatial details (Le Grand et al., 2001, 2004; Vogelsang et al., 2018).

In comparison, we found that CNNs trained with a sequence of blurry to clear objects led to a much more specialized form of learning. After each stage of training, performance was selectively elevated at the most recently trained blur level, whereas generalization to neighboring blur levels was extremely poor. This pattern of results replicated in CNNs trained on subordinate-level classification of dogs and birds, as well as a CNN trained to distinguish animate from inanimate objects. These changes in test performance could be explained by the progressive shift in the network’s preference for higher spatial frequency information to leverage better performance. Our findings suggest that object-trained CNNs are predisposed to learn specialized representations that are highly specific in spatial scale, such that changes in the spatial frequency content of the training images cause previously learned representations to be modified or overwritten. To the extent that CNNs provide a suitable model of the learned representations of the human visual system, these findings lead us to conclude that poor acuity in infancy is unlikely to account for the robustness with which people can recognize blurry objects in adulthood (e.g., **Figure 1B**).

Perhaps more importantly, our findings demonstrate how faces and objects are processed in a qualitatively different manner by CNNs when they are trained using a common progression of blurry to clear images. We observed a growing divergence in the spatial tuning of face- and object-trained CNNs after each stage of blur training. These results provide novel computational evidence to suggest that faces constitute a special stimulus class that can be processed more holistically than most other object categories, by utilizing information in lower spatial frequency bands for successful recognition.

Our findings add to a large body of studies of holistic processing, which have relied on a variety of experimental methods and operational definitions to characterize this process (Goffaux & Rossion, 2006; Le Grand et al., 2004; Michel, Rossion, Han, Chung, & Caldara, 2006; Richler & Gauthier, 2014; Richler, Gauthier, Wenger, & Palmeri, 2008; Tanaka & Farah, 1993; Tanaka & Simonyi, 2016), and lead us to suggest a possible computational definition of this inferred process. Specifically, holistic processing may entail the ability to recognize a set of stimuli across a range of blur levels while relying on a common set of visual representations. An important component of face recognition involves distinguishing subtle differences in the overall 3D shape of different faces (Chang & Tsao, 2017), and shape from shading provides a critical cue for encoding information about face shape (Atick, Griffin, & Redlich, 1996; Yildirim, Belledonne, Freiwald, & Tenenbaum, 2020). These cues about 3D face shape take the form of gradual rather than abrupt changes in luminance that occur across the 2D facial image. The gradual nature of these luminance changes may partly explain why face recognition can tolerate fairly high levels of blur (**Figure 1A**).

Consistent with this notion, previous research has suggested that face recognition tends to rely on information in the lower and mid-range of spatial frequencies. Optimal levels of face recognition performance have been reported for bandpass filtered faces typically centered at around 8 cycles per stimulus (Gaspar et al., 2008; Näsänen, 1999; Peli et al., 1994). Another study found that low-pass filtered faces with hair can still be successfully recognized at a cut-off frequency of only 2.6 cycles per stimulus (Kwon & Legge, 2011), while those without hair can be recognized at a cut-off frequency of 4.2 cycles per stimulus. (It should be noted that letter identification was successful with even lower cut-off frequencies of ~1 cycle per stimulus.) There is also some evidence to suggest that faces can be recognized at lower spatial frequencies than certain non-face objects, such as airplanes (Harel & Bentin, 2009). Although such findings point to potential differences regarding how faces and objects are processed across varying spatial scales, the findings reported here indicate a much sharper divergence between face and object processing.

Our computational findings suggest that the robust recognition of blurry objects cannot readily be attributed to the blurry nature of early infant vision. How then is such robustness achieved and maintained in adulthood? We propose that visual experience at all ages is permeated with experiences of blurry objects, such as whenever they appear widely separated in depth from the point of fixation (Sprague et al., 2016). There is also evidence to suggest that whenever an observer fixates an object at a new depth plane, the accommodation system requires about 200ms to adjust its focus (Chirre, Prieto, & Artal, 2015). It is also the case that vision in the periphery leads to frequent sampling of low resolution objects, and that the object recognition system may be incentivized to learn the appearance of these peripheral objects to guide the process of visual search (Li & DiCarlo, 2008; Strasburger et al., 2011). As a consequence, the human visual system experiences continual training with a wide range of blur levels in the real world. It will be of interest for future studies to investigate whether CNNs, trained with different regimes of blurry experiences, can acquire robustness to blurry objects in a manner that better matches human performance. In addition to blur, there is growing interest in understanding why CNNs lack robustness to challenging viewing conditions such as visual noise, occlusion, and other types of image distortion (Geirhos et al., 2018; H. Jang & Tong, 2018; Spoerer, McClure, & Kriegeskorte, 2017). Although CNNs can be trained to become more robust to a specific type of image distortion, such training typically fails to generalize to other challenging visual conditions. A major goal in computer vision is to develop deep neural networks that can learn to robustly recognize objects from fewer training examples to better mimic human vision.

Although we found that blurry to clear image training was not effective for promoting robustness of object recognition, the concept of coarse-to-fine processing has been widely discussed in the object recognition literature (Bullier, 2001; Schyns & Oliva, 1994; Watt, 1987). By some accounts, low spatial frequency information processing serves to guide the processing of high spatial frequency information. Although this proposal does not necessarily justify our strategy of training CNNs with blurry to clear images, such a training regime could provide, in some instances, computational advantages in learning. From an optimization perspective, initial training with blurry inputs could serve to flatten the parameter search space and act as a form of regularization, thereby promoting more efficient search for a global minimum.

Finally, this study demonstrates the utility of using deep neural networks to address questions pertaining to human visual development, which may otherwise be difficult or impossible to test. Although one can observe or record the visual experiences of infants, it is rarely possible to modify their experiences outside of a brief experimental session. By contrast, the totality of a deep network’s learned experiences can be precisely specified and manipulated, and any control conditions of interest can also be tested. Thus, the impact of a series of learning experiences, from beginning to end, can be studied in detail. For example, here we could evaluate whether blurry to clear image training might be beneficial for learning distinctions between animate and inanimate objects, as developmental research has suggested that categorization ability gradually progresses from superordinate to basic to subordinate levels (Mandler & McDonough, 1993; Quinn, 2004). Although we did not find any benefit of blurry to clear image training with respect to acquiring robustness to blur, it is conceivable that future studies could investigate other questions pertaining to level of categorization and visual development. There is also growing interest in using ecologically valid input to train deep networks, such as obtaining video data from babies using head-mounted cameras (Smith, Jayaraman, Clerkin, & Yu, 2018). For example, a recent study demonstrated that a contrastive method of unsupervised learning allowed a deep neural network to learn appropriate object representations from baby-cam data (Zhuang et al., 2021). This confluence of neuroscience, vision science, and artificial intelligence is progressing rapidly (Cichy & Kaiser, 2019; Kar, Kubilius, Schmidt, Issa, & DiCarlo, 2019; Kriegeskorte, 2015; Yamins & DiCarlo, 2016). Soon, through the development of more biologically plausible network architectures (e.g., Kar et al., 2019; Kietzmann et al., 2019), better conceived learning methods (e.g, Zhuang et al., 2021), and the use of ecologically and developmentally valid training data, neuroscientists will be well-positioned to develop better models of the human visual system.

## Acknowledgements

We would like to thank Kaylee Bashor for technical support. This research was supported by a grant from the National Eye Institute (R01EY029278) to FT, and a core grant from the National Eye Institute (P30-EY008126) to the Vanderbilt Vision Research Center (Director David Calkins).

## Author Contributions

HJ and FT devised and designed the study; HJ coded the experiments, CNNs, and performed all data analyses; HJ and FT wrote the paper together.

## References

Atick, J. J., Griffin, P. A., & Redlich, A. N. (1996). Statistical approach to shape from shading: reconstruction of three-dimensional face surfaces from single two-dimensional images. Neural Comput, 8(6), 1321–1340.

Boucart, M., Lenoble, Q., Quettelart, J., Szaffarczyk, S., Despretz, P., & Thorpe, S. J. (2016). Finding faces, animals, and vehicles in far peripheral vision. Journal of Vision, 16(2), 10–10.

Braddick, O., & Atkinson, J. (2011). Development of human visual function. Vision Research, 51(13), 1588–1609.

Bullier, J. (2001). Integrated model of visual processing. Brain Research Reviews, 36(2-3), 96–107.

Chang, L., & Tsao, D. Y. (2017). The Code for Facial Identity in the Primate Brain. Cell, 169(6), 1013–1028 e1014.

Chirre, E., Prieto, P., & Artal, P. (2015). Dynamics of the near response under natural viewing conditions with an open-view sensor. Biomed Opt Express, 6(10), 4200–4211.

Cichy, R. M., & Kaiser, D. (2019). Deep Neural Networks as Scientific Models. Trends Cogn Sci, 23(4), 305–317.

Diamond, R., & Carey, S. (1986). Why faces are and are not special: an effect of expertise. J Exp Psychol Gen, 115(2), 107–117.

Dobson, V., & Teller, D. Y. (1978). Visual acuity in human infants: a review and comparison of behavioral and electrophysiological studies. Vision Research, 18(11), 1469–1483.

Dodge, S., & Karam, L. (2017). A study and comparison of human and deep learning recognition performance under visual distortions. Paper presented at the 2017 26th International Conference on Computer Communication and Networks.

Duncan, R. O., & Boynton, G. M. (2003). Cortical magnification within human primary visual cortex correlates with acuity thresholds. Neuron, 38(4), 659–671.

Freiwald, W. A., Tsao, D. Y., & Livingstone, M. S. (2009). A face feature space in the macaque temporal lobe. Nat Neurosci, 12(9), 1187–1196.

Fukushima, K. (1980). Neocognitron: A self-organizing neural network model for a mechanism of pattern recognition unaffected by shift in position. Biological Cybernetics, 36(4), 193–202.

Gaspar, C., Sekuler, A. B., & Bennett, P. J. (2008). Spatial frequency tuning of upright and inverted face identification. Vision Res, 48(28), 2817–2826.

Gauthier, I., & Tarr, M. J. (1997). Becoming a "Greeble" expert: exploring mechanisms for face recognition. Vision Res, 37(12), 1673–1682.

Geirhos, R., Medina Temme, C. R., Rauber, J., Schutt, H. H., Bethge, M., & Wichmann, F. A. (2018). Generalisation in humans and deep neural networks. Paper presented at the Neural Information Processing Systems.

Geldart, S., Mondloch, C. J., Maurer, D., De Schonen, S., & Brent, H. P. (2002). The effect of early visual deprivation on the development of face processing. Developmental Science, 5(4), 490–501.

Goffaux, V., Hault, B., Michel, C., Vuong, Q. C., & Rossion, B. (2005). The respective role of low and high spatial frequencies in supporting configural and featural processing of faces. Perception, 34(1), 77–86.

Goffaux, V., & Rossion, B. (2006). Faces are “spatial”--holistic face perception is supported by low spatial frequencies. Journal of Experimental Psychology: Human Perception and Performance, 32(4), 1023.

Goodfellow, I. J., Shlens, J., & Szegedy, C. (2014). Explaining and harnessing adversarial examples. [Electronic Version]. arXiv.

Güçlü, U., & van Gerven, M. A. (2015). Deep neural networks reveal a gradient in the complexity of neural representations across the ventral stream. Journal of Neuroscience, 35(27), 10005–10014.

Harel, A., & Bentin, S. (2009). Stimulus type, level of categorization, and spatial-frequencies utilization: implications for perceptual categorization hierarchies. Journal of Experimental Psychology: Human Perception and Performance, 35(4), 1264.

He, K., Zhang, X., Ren, S., & Sun, J. (2015). Delving deep into rectifiers: Surpassing human-level performance on imagenet classification. Paper presented at the Proceedings of the IEEE International Conference on Computer Vision.

Hebart, M. N., Zheng, C. Y., Pereira, F., & Baker, C. I. (2020). Revealing the multidimensional mental representations of natural objects underlying human similarity judgements. Nat Hum Behav, 4(11), 1173–1185.

Horikawa, T., & Kamitani, Y. (2017). Generic decoding of seen and imagined objects using hierarchical visual features. Nat Commun, 8, 15037.

Hubel, D. H., & Wiesel, T. N. (1962). Receptive fields, binocular interaction and functional architecture in the cat’s visual cortex.. Journal of Physiology, 160, 106–154.

Jang, H., McCormack, D., & Tong, F. (2017). Evaluating the robustness of object recognition to visual noise in humans and convolutional neural networks. Journal of Vision, 17(10), 805.

Jang, H., & Tong, F. (2018). Can deep learning networks acquire the robustness of human recognition when faced with objects in visual noise? Paper presented at the Vision Sciences Society.

Kanwisher, N., McDermott, J., & Chun, M. M. (1997). The fusiform face area: a module in human extrastriate cortex specialized for face perception. Journal of Neuroscience, 17(11), 4302–4311.

Kar, K., Kubilius, J., Schmidt, K., Issa, E. B., & DiCarlo, J. J. (2019). Evidence that recurrent circuits are critical to the ventral stream’s execution of core object recognition behavior. Nature Neuroscience, 22(6), 974–983.

Khaligh-Razavi, S. M., & Kriegeskorte, N. (2014). Deep supervised, but not unsupervised, models may explain IT cortical representation. PLoS Comput Biol, 10(11), e1003915.

Kietzmann, T. C., Spoerer, C. J., Sorensen, L. K. A., Cichy, R. M., Hauk, O., & Kriegeskorte, N. (2019). Recurrence is required to capture the representational dynamics of the human visual system. Proc Natl Acad Sci U S A, 116(43), 21854–21863.

Kiorpes, L., & Movshon, J. A. (2004). Neural limitations on visual development in primates. The visual neurosciences, 1, 159–173.

Kriegeskorte, N. (2015). Deep Neural Networks: A New Framework for Modeling Biological Vision and Brain Information Processing. Annu Rev Vis Sci, 1, 417–446.

Krizhevsky, A., Sutskever, I., & Hinton, G. E. (2012). Imagenet classification with deep convolutional neural networks. Paper presented at the Advances in Neural Information Processing Systems.

Kwon, M., & Legge, G. E. (2011). Spatial-frequency cutoff requirements for pattern recognition in central and peripheral vision. Vision Res, 51(18), 1995–2007.

Le Grand, R., Mondloch, C. J., Maurer, D., & Brent, H. P. (2001). Early visual experience and face processing. Nature, 410(6831), 890–890.

Le Grand, R., Mondloch, C. J., Maurer, D., & Brent, H. P. (2004). Impairment in holistic face processing following early visual deprivation. Psychol Sci, 15(11), 762–768.

LeCun, Y., Bengio, Y., & Hinton, G. (2015). Deep learning. Nature, 521(7553), 436–444.

Lewis, T. L., Ellemberg, D., Maurer, D., Wilkinson, F., Wilson, H. R., Dirks, M., et al. (2002). Sensitivity to global form in glass patterns after early visual deprivation in humans. Vision Res, 42(8), 939–948.

Li, N., & DiCarlo, J. J. (2008). Unsupervised natural experience rapidly alters invariant object representation in visual cortex. Science, 321(5895), 1502–1507.

Loffler, G., Yourganov, G., Wilkinson, F., & Wilson, H. R. (2005). fMRI evidence for the neural representation of faces. Nat Neurosci, 8(10), 1386–1390.

Mandler, J. M., & McDonough, L. (1993). Concept formation in infancy. Child Development, 8, 291–318.

Michel, C., Rossion, B., Han, J., Chung, C. S., & Caldara, R. (2006). Holistic processing is finely tuned for faces of one’s own race. Psychol Sci, 17(7), 608–615.

Näsänen, R. (1999). Spatial frequency bandwidth used in the recognition of facial images. Vision Research, 39(23), 3824–3833.

Ng, H. W., & Winkler, S. (2014). A data-driven approach to cleaning large face datasets. Paper presented at the 2014 IEEE International Conference on Image Processing.

Norcia, A. M., & Tyler, C. W. (1985). Spatial frequency sweep VEP: visual acuity during the first year of life. Vision Res, 25(10), 1399–1408.

Peli, E., Lee, E., Trempe, C. L., & Buzney, S. (1994). Image enhancement for the visually impaired: the effects of enhancement on face recognition. J Opt Soc Am A Opt Image Sci Vis, 11(7), 1929–1939.

Phillips, P. J., Yates, A. N., Hu, Y., Hahn, C. A., Noyes, E., Jackson, K., et al. (2018). Face recognition accuracy of forensic examiners, superrecognizers, and face recognition algorithms. Proceedings of the National Academy of Sciences, 115(24), 6171–6176.

Quinn, P. C. (2004). Development of subordinate-level categorization in 3-to 7-month-old infants. Child Dev, 75(3), 886–899.

Richler, J. J., & Gauthier, I. (2014). A meta-analysis and review of holistic face processing. Psychol Bull, 140(5), 1281–1302.

Richler, J. J., Gauthier, I., Wenger, M. J., & Palmeri, T. J. (2008). Holistic processing of faces: perceptual and decisional components. J Exp Psychol Learn Mem Cogn, 34(2), 328–342.

Russakovsky, O., Deng, J., Su, H., Krause, J., Satheesh, S., Ma, S., et al. (2015). Imagenet large scale visual recognition challenge. International journal of computer vision, 115(3), 211–252.

Schyns, P. G., & Oliva, A. (1994). From blobs to boundary edges: Evidence for time-and spatial-scale-dependent scene recognition. Psychological Science, 5(4), 195–200.

Simonyan, K., & Zisserman, A. (2014). Very deep convolutional networks for large-scale image recognition. arXiv preprint arXiv:1409.1556.

Sinha, P. (2002). Recognizing complex patterns. Nature Neuroscience, 5(11), 1093–1097.

Smith, L. B., Jayaraman, S., Clerkin, E., & Yu, C. (2018). The Developing Infant Creates a Curriculum for Statistical Learning. Trends Cogn Sci, 22(4), 325–336.

Spoerer, C. J., McClure, P., & Kriegeskorte, N. (2017). Recurrent Convolutional Neural Networks: A Better Model of Biological Object Recognition. Front Psychol, 8, 1551.

Sprague, W. W., Cooper, E. A., Reissier, S., Yellapragada, B., & Banks, M. S. (2016). The natural statistics of blur. J Vis, 16(10), 23.

Strasburger, H., Rentschler, I., & Jüttner, M. (2011). Peripheral vision and pattern recognition: A review. Journal of Vision, 11(5), 13–13.

Taigman, Y., Yang, M., Ranzato, M. A., & Wolf, L. (2014). Deepface: Closing the gap to human-level performance in face verification. Paper presented at the Proceedings of the IEEE conference on Computer Vision and Pattern Recognition.

Tanaka, J. W., & Farah, M. J. (1993). Parts and wholes in face recognition. Q J Exp Psychol A, 46(2), 225–245.

Tanaka, J. W., & Simonyi, D. (2016). The “parts and wholes” of face recognition: A review of the literature. Q J Exp Psychol (Hove), 69(10), 1876–1889.

Thorpe, S. J., Gegenfurtner, K. R., Fabre‐Thorpe, M., & Bülthoff, H. H. (2001). Detection of animals in natural images using far peripheral vision. European Journal of Neuroscience, 14(5), 869–876.

Tong, F., Nakayama, K., Moscovitch, M., Weinrib, O., & Kanwisher, N. (2000). Response properties of the human fusiform face area. Cognitive Neuropsychology, 17, 257–279.

Virsu, V., & Rovamo, J. (1979). Visual resolution, contrast sensitivity, and the cortical magnification factor. Exp Brain Res, 37(3), 475–494.

Vogelsang, L., Gilad-Gutnick, S., Ehrenberg, E., Yonas, A., Diamond, S., Held, R., et al. (2018). Potential downside of high initial visual acuity. Proceedings of the National Academy of Sciences, 115(44), 11333–11338.

Watt, R. (1987). Scanning from coarse to fine spatial scales in the human visual system after the onset of a stimulus. J Opt Soc Am A, 4(10), 2006–2021.

Wood, I. C., Hodi, S., & Morgan, L. (1995). Longitudinal change of refractive error in infants during the first year of life. Eye (Lond), 9 (Pt 5), 551–557.

Yamins, D. L., & DiCarlo, J. J. (2016). Using goal-driven deep learning models to understand sensory cortex. Nat Neurosci, 19(3), 356–365.

Yamins, D. L., Hong, H., Cadieu, C. F., Solomon, E. A., Seibert, D., & DiCarlo, J. J. (2014). Performance-optimized hierarchical models predict neural responses in higher visual cortex. Proc Natl Acad Sci U S A, 111(23), 8619–8624.

Yildirim, I., Belledonne, M., Freiwald, W., & Tenenbaum, J. (2020). Efficient inverse graphics in biological face processing. Sci Adv, 6(10), eaax5979.

Zhuang, C., Yan, S., Nayebi, A., Schrimpf, M., Frank, M. C., DiCarlo, J. J., et al. (2021). Unsupervised neural network models of the ventral visual stream. Proc Natl Acad Sci U S A, 118(3).

