## Supplementary Figure for "Convolutional neural networks trained with a developmental sequence of blurry to clear images reveal core differences between face and object processing"

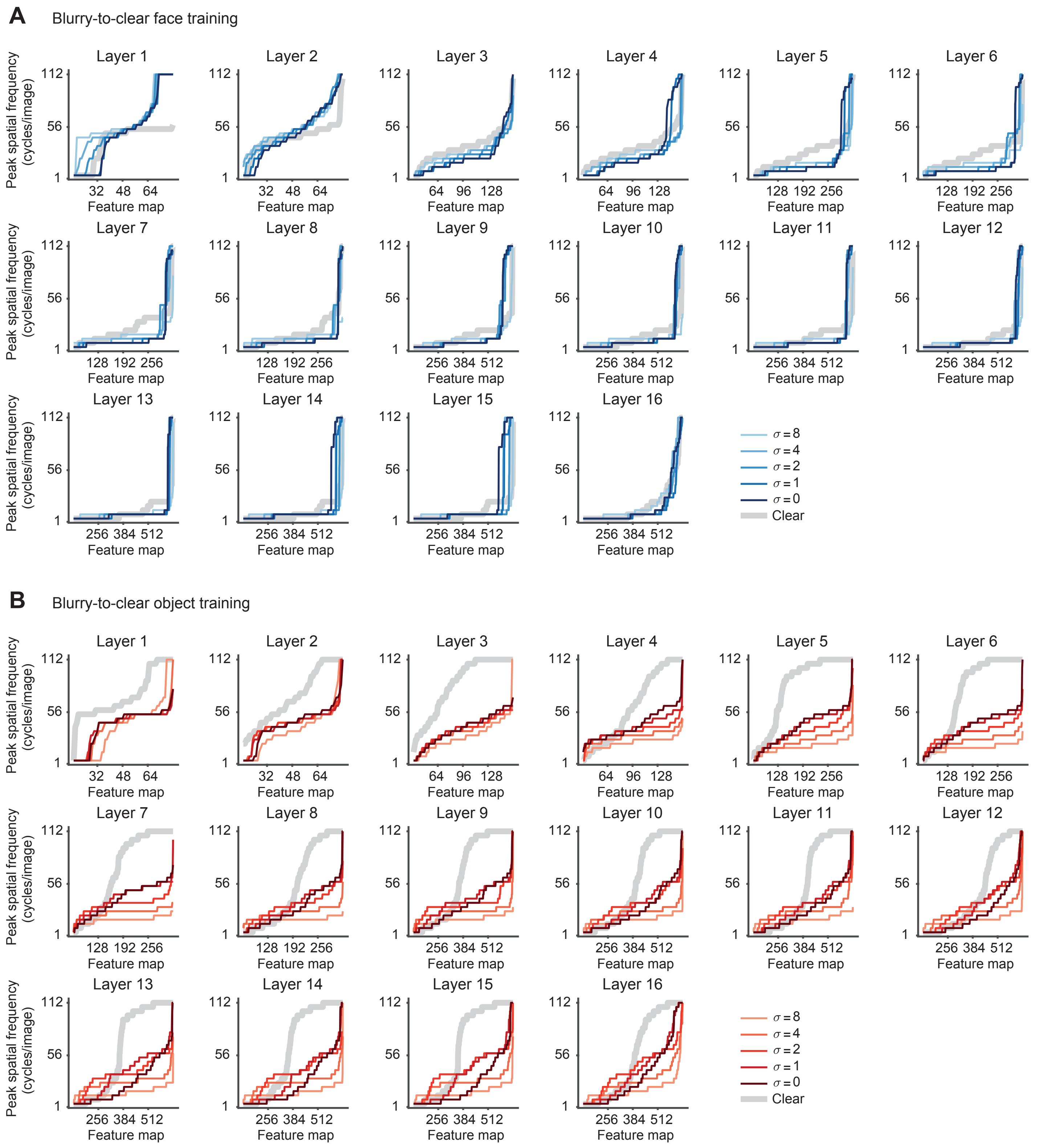


**Supplementary Figure 1.** Spatial frequency preferences of VGG-19, following the procedures as in **Figure 4C-D**. Peak spatial frequency preferences of the face-trained (**A**) and object-trained (**B**) networks are shown. Note that VGG19 uses small 3×3 convolutional filters in all convolutional layers, so the first few layers have very small receptive fields and allow for little variation in spatial frequency preference.
